# NERD-seq: A novel approach of Nanopore direct RNA sequencing that expands representation of non-coding RNAs

**DOI:** 10.1101/2021.05.06.442990

**Authors:** Luke Saville, Yubo Cheng, Babita Gollen, Liam Mitchell, Matthew Stuart-Edwards, Travis Haight, Majid Mohajerani, Athanasios Zovoilis

## Abstract

The new next-generation sequencing platforms by Oxford Nanopore Technologies for direct RNA sequencing (direct RNA-seq) allow for an in-depth and comprehensive study of the epitranscriptome by enabling direct base calling of RNA modifications. Non-coding RNAs constitute the most frequently documented targets for RNA modifications. However, the current standard direct RNA-seq approach is unable to detect many of these RNAs. Here we present NERD-seq, a sequencing approach which enables the detection of multiple classes of non-coding RNAs excluded by the current standard approach. Using total RNA from a tissue with high known transcriptional and non-coding RNA activity in mouse, the brain hippocampus, we show that, in addition to detecting polyadenylated coding and non-coding transcripts as the standard approach does, NERD-seq is able to significantly expand the representation for other classes of RNAs such as snoRNAs, snRNAs, scRNAs, srpRNAs, tRNAs, rRFs and non-coding RNAs originating from LINE L1 elements. Thus, NERD-seq presents a new comprehensive direct RNA-seq approach for the study of epitranscriptomes in brain tissues and beyond.

## INTRODUCTION

Since the advent of RNA sequencing (1,2), our understanding of the transcriptome and its regulation has grown substantially(3). By improving the context of transcriptome analysis, RNA sequencing allows for an improved understanding of the effects of cellular regulation(4,5), the environment(6,7), and disease pathology(8,9), on transcription changes and regulation.

Since RNA sequencing’s introduction as an essential molecular biology tool, many technological iterations have occurred. From the initial iteration of sequencing by synthesis of cDNA, through improved technologies such as 454 pyrosequencing(10), and eventually the Illumina sequencing platform(11), a number of limitations have been identified. In particular, although short read sequencing is both powerful and exceptionally high throughput on the Illumina sequencing platform, the need to fragment longer RNA polynucleotides and the need to reverse transcribe the RNA and PCR amplify the cDNA before the sequencing makes identification of RNA modifications(12), and variations in RNA structure and RNA splicing(13,14) challenging, complicated and expensive, while increasing the probability of sequencing and analysis errors(15–17).

Recent technological iterations show promise in addressing many of the above challenges associated with RNA sequencing. Third-generation sequencing is dominated by the PacBio and Oxford Nanopore Technologies (ONT) sequencing platforms and has made long read – whole molecule – polynucleotide sequencing possible(18–22). Most promising, the Nanopore platforms enable us to sequence native RNA polynucleotides in their whole form, without the need for replacement by cDNA and subsequent amplification(23–25). Although Nanopore sequencing is capable of sequencing cDNA and cDNA amplicons(25), the sequencing of the native RNA strand (direct RNA-seq) allows the resolution of modified RNA nucleotides to provide context to the epitranscriptome(19,26), while reducing the library preparation and analysis complexity(27,28).

Over 100 unique RNA modifications have been described(29), including adenosine to inosine (A-to-I) edits, pseudouridylation, and methylation on multiple sites on the nucleotide base(30–35). Non-coding RNAs (ncRNAs) constitute frequent targets of these modifications with emerging significance for human health and disease. For example, higher A-to-I editing ratios in SINE RNAs have been linked to reduced severity in some viral infections such as that by SARS-CoV-2(36), and reduced A-to-I editing ratios have been linked to multiple sclerosis(37). Additionally, m6A modifications are thought to encourage circular RNA formation by back splicing(38) and inosines in tRNAs allow for wobble base pairing for redundant codon recognition, dysfunction of which may cause intellectual disability(39). Direct RNA-seq has exhibited the ability to detect these RNA modifications in recent research studies(28,40,41) by analyzing perturbations in the ionic trace plots produced by Nanopore sequencing.

Despite the promise of Nanopore sequencing to deconvolute the above native edits, the process of direct RNA-seq using standard, polyadenylation (poly(A))-selecting Nanopore library preparations, eliminates the possibility for capturing many of the ncRNAs that constitute the vast majority of editing substrates without modifying the library preparation method(23,32,42–44). Some of the substrates that may be omitted include among others tRNAs, snoRNAs, snRNAs, scRNAs and other cellular non-poly(A) RNAs as well as viral RNAs. Here, we present an approach, called NERD-seq, to expand the ncRNA representation in Nanopore direct RNA-seq to include these classes of ncRNAs.

## MATERIALS AND METHODS

### Post mortem hippocampal tissues

Mice were raised and had tissue extracted as described previously(45). Left and right mouse hippocampus tissue were homogenized separately in 1.0mL Trizol reagent: 15-minute incubation and subsequent grinding using a pestle until nothing but insoluble connective tissue remained. The homogenized mix was pipetted up and down and the solution was stored at −80°C. 0.5mL of homogenized mixture was phase separate by the addition of 100μL of chloroform (Sigma, C2432) and mixed by inversion, incubated for three minutes and centrifuged at 12,500xg for fifteen minutes at 4°C. The top (aqueous) layer was transferred to a new tube and mixed with 250μL of isopropanol (Fisher, 67-63-0), followed by a 1-hour incubation at −20°C and centrifugation at 12,500xg for ten minutes at 4°C. The supernatant was removed and the pellet was washed and mixed with 0.5mL of 75% ethanol, followed by a centrifugation at 7 600xg for five minutes at 4°C. The supernatant was removed and the pellet was allowed to dry for 1 minute before eluting in 30μL of nuclease free H_2_O. The eluted RNA was heated at 55°C for fifteen minutes, and subsequently incubated with 1μL of DNaseI (NEB, M0303), 10μL of 10x DNaseI buffer (NEB, B0303), and 39μL of nuclease free H_2_O for 15 minutes at 37°C. The RNA was further cleaned using the Zymo Research RNA clean and concentrator kit −25 (R1017) and combined in equal densities (left and right RNA) The RNA was stored at −80°C.

### Direct RNA sequencing using Nanopore

For the standard approach we used the ONT SQK_RNA002 kit and the direct RNA-seq protocol, listed at the Nanopore Community portal (https://nanoporetech.com/community) with 1.5ug total RNA as starting material. In both the standard and the the NRED-seq protocol, ligations were performed for 15min instead of 10min, to increase yield. For NERD-seq, the standard protocol was further modified as follows: 1.5ug total RNA was separated into two fractions (short and long RNA) using the Invitrogen MirVana kit (AM1561) with a modified protocol as described before in our short RNA-seq approach for illumina (45–48). 90uL of the short RNA elution fraction was polyadenylated using 12uL 10x polyA polymerase buffer (NEB, B0276), 12uL 10mM ATP (NEB, B0756A), 6uL polyA polymerase (NEB, M0276) and incubated at 37C for 30min. The polyadenylated short fraction and the long fraction were combined and purified and concentrated using the RNeasy MinElute kit (Qiagen, 74204). 12uL of the RNA was adaptor ligated using 4uL NEBNext Quick ligation buffer (B6058), 0.66uL RNA CS (Nanopore), 1.3uL RTA adaptor (Nanopore), 2uL 2 000 000U T4 DNA ligase (NEB, M0202). The reaction was incubated at room temperature for 10 minutes. The sample was subsequently reverse transcribed using 1.5uL of the Lucigen OmniAmp polymerase (F831942-1), 10uL 10x OmniAmp buffer (Lucigen, F883707-1), 8uL dNTP (NEB, N0447), 6uL 100mM MgSO_4_ (Lucigen, F98695-1), 10uL Betaine (Lucigen, F881901-1), 5uL random primer mix (NEB, S1330), and 39.5uL RNase free H_2_O. The mixture was incubated for 50C – 10min, 70C – 20min. Samples were cleaned by 2.88x Omega Mag-Bind^®^ TotalPure NGS beads. The 20uL library was ligated to the Nanopore RMX adaptor using 6uL RMX adaptor (Nanopore), 8uL NEBNext quick ligation buffer, 3uL RNase free H_2_O, and 3uL 2 000 000U T4 DNA ligase and incubated for 10 minutes at room temperature. The adapted library was 1x bead cleaned using the Omega Mag-Bind^®^ TotalPure NGS beads, eluted in Nanopore elution buffer. Library was loaded onto Nanopore PromethION according to manufacturer’s instructions (flow cells were version 9.4.1).

### Bioinformatics analysis

The RNA sequencing data was aligned using minimap2 (PMID: 29750242) version 2.17 against the GRCm38/mm10 Mouse Genome. Minimap2 was used with the options: -x sr to optimize for short RNA reads. Aligned sam files were converted to bam format using samtools version 1.7(49). Bam to bed conversion was performed using bedtools version v2.26.0(50). Intersections were performed using bedtools v2.26.0 and with the intersect options: -s -wa -u -e -f 0.8. Annotations were retrieved from UCSC Table Browser (https://genome.ucsc.edu/cgi-bin/hgTables); specifically, GENCODE V23, GENCODE V25, and RepeatMasker (as of Jan 2021); with subsequent filtering for specific families of elements. The number of lines in the intersected bed files (corresponding to number of reads which occur within respective annotation file) are taken as a percentage of the total number of reads present in the pre-intersection bed file, thus giving proportion of reads which fall into the category represented by each annotation. TSS annotation is based on the Eponine annotation (51), and rRNA annotation was done using the Seqmonk default annotation. Models of read distribution around the TSS of various genomic elements were performed using the Babraham NGS analysis suite Seqmonk 1.38.2 (https://www.bioinformatics.babraham.ac.uk/projects/seqmonk/). In brief, we constructed metagene models around a hypothetical set of genomic points, such as the transcription start site and subsequently plotted the distribution of read counts from all contributing elements for each position. Then the numbers of reads at the same strand with the element around each different TSS were calculated and attributed to defined points in the model. Relative density or cumulative distributions at the metagene plots were generated using Seqmonk.

### Data access

NERD-seq data have been deposited to GEO with access number GSE171312 (RNA sample used in the current study is named as “ctrl-6m_3”).

The external dataset from Sessegolo *et al*, 2019 was accessed from the European Nucleotide Archive, accession: PRJEB27590 (https://www.genoscope.cns.fr/ont_mouse_rna/datasets_RNA_LR.html). Samples Brain C1 and C2 were used for comparison.

## RESULTS

### The modified library construction protocol used in NERD-seq bypasses the limitations posed by the standard Nanopore direct RNA-seq approach

Standard, direct RNA-seq relies on poly(A) selection by using a poly(T)-tethering adaptor that base-pairs canonically to the motor protein linked adaptor, facilitating the movement of the polynucleotide string through the protein pore (Figure 1A). Nucleotide detection is performed by analyzing perturbations of the ionic current in the sequencing membrane(25). By doing so, Nanopore sequencing provides a moderately simple method to sequence the poly(A) transcriptome, which predominantly represents post-transcriptionally modified mRNAs(23). Unfortunately, this method omits many ncRNAs that do not have a poly(A) tail.

**Figure 1.**
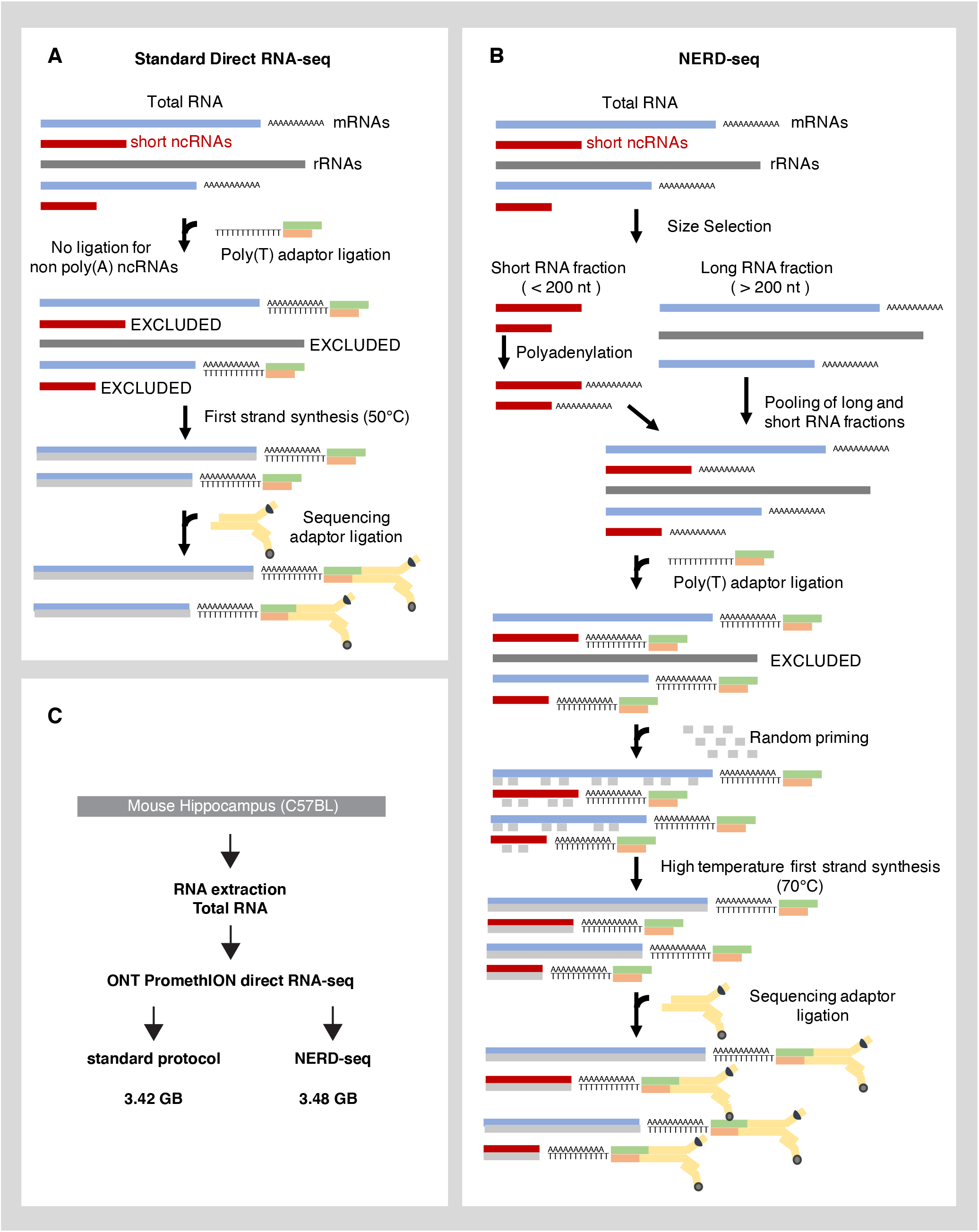
Description of the NERD-seq approach. **(A)** Illustration of the standard Nanopore direct RNA-seq approach and the types of RNAs it can theoretically detect. ncRNA: non-coding RNA; rRNA: ribosomal RNA; mRNA: messenger RNA **(B)** Illustration of the NERD-seq approach and the additional types of RNAs (short non poly(A) ncRNAs) it can theoretically detect. **(C)** Experimental design of the current study.

To address this, we established a Non-coding Enriched RNA Direct sequencing approach called NERD-seq. In this modified approach (summarized in Figure 1B), we first separated total RNA into two fractions (long and short RNA fraction) using a column-based size enrichment approach, which we have used in the past for similar approaches to enrich for short (approx. <200nt) ncRNAs with Illumina sequencing (short-RNA-seq)(45,48,52). This column-based approach enables for the significant enrichment of short transcripts (although it does not completely eliminate very long RNAs)(53). To the short fraction, we added a polyadenine tail using a poly(A) polymerase to allow in subsequent steps the short ncRNAs to form canonical base-pairing to the poly(T) sequencing adaptor and, thus, facilitate its ligation. Additionally, this poly(A) addition helps achieve higher accuracy in Nanopore sequencing of short RNAs because the beginning of the sequence usually requires some voltage adjustment for accurate basecalling and, proportionally, the error rate becomes higher for short reads(54,55). At the same time, full length rRNAs that remain at the long RNA fraction, and are usually not desirable due to their extremely high read numbers that decrease yield for the other RNA classes, are not subjected to the polyadenylation, enabling their exclusion at later protocol steps. After deactivating the poly(A) polymerase, the two fractions were pooled again together. Thus, the new sample contains all the initial RNAs with a fraction of the short ones being poly-adenylated.

An additional problem that arises in both Illumina and Nanopore sequencing protocols is the ability of highly structured RNA regions to hinder the sequencing process in various ways. In the case of Nanopore sequencing, highly structured RNA regions may prevent the pulling of the RNA molecule through the pore(25). To this end, although the direct RNA-seq protocol does not sequence cDNA, an optional first-strand cDNA synthesis step through a reverse transcriptase is recommended to stabilize the native RNA strand and resolve highly structured RNA regions commonly observed in SINE RNAs, snoRNAs and other ncRNAs (the cDNA strand is not sequenced)(56–58). However, the direct RNA-seq protocol utilizes reverse transcriptases with a temperature optimum between 45°C-50°C. At this temperature range some highly structured RNA regions remain unresolved resulting in reverse transcriptase pauses and subsequent decrease in the length of sequenced transcripts (Figure 2A, grey distribution). To this end, we modified the complementary strand synthesis in two ways: Firstly, we reverse transcribed using the OmniAmp polymerase (Lucigen, F831942-1). Omniamp is a mutated form of the PyroPhage 3173 DNA polymerase(59,60). Omniamp is capable of loop-mediated isothermal amplification that enables reverse transcription at a temperature as high as 70°C, which facilitates further RNA unfolding. Moreover, to prevent RNA degradation at this temperature, we added random primers and initially allowed first-strand synthesis by OmniAmp for 10 minutes at 50°C to “coat” RNAs with protective cDNA strand fragments in multiple locations along their whole length. Since OmniAmp has a strong strand displacement activity(56), all these short first-strand fragments are subsequently displaced by the reverse transcribed cDNA strand that is initiated by the poly(T) adaptor.

**Figure 2.**
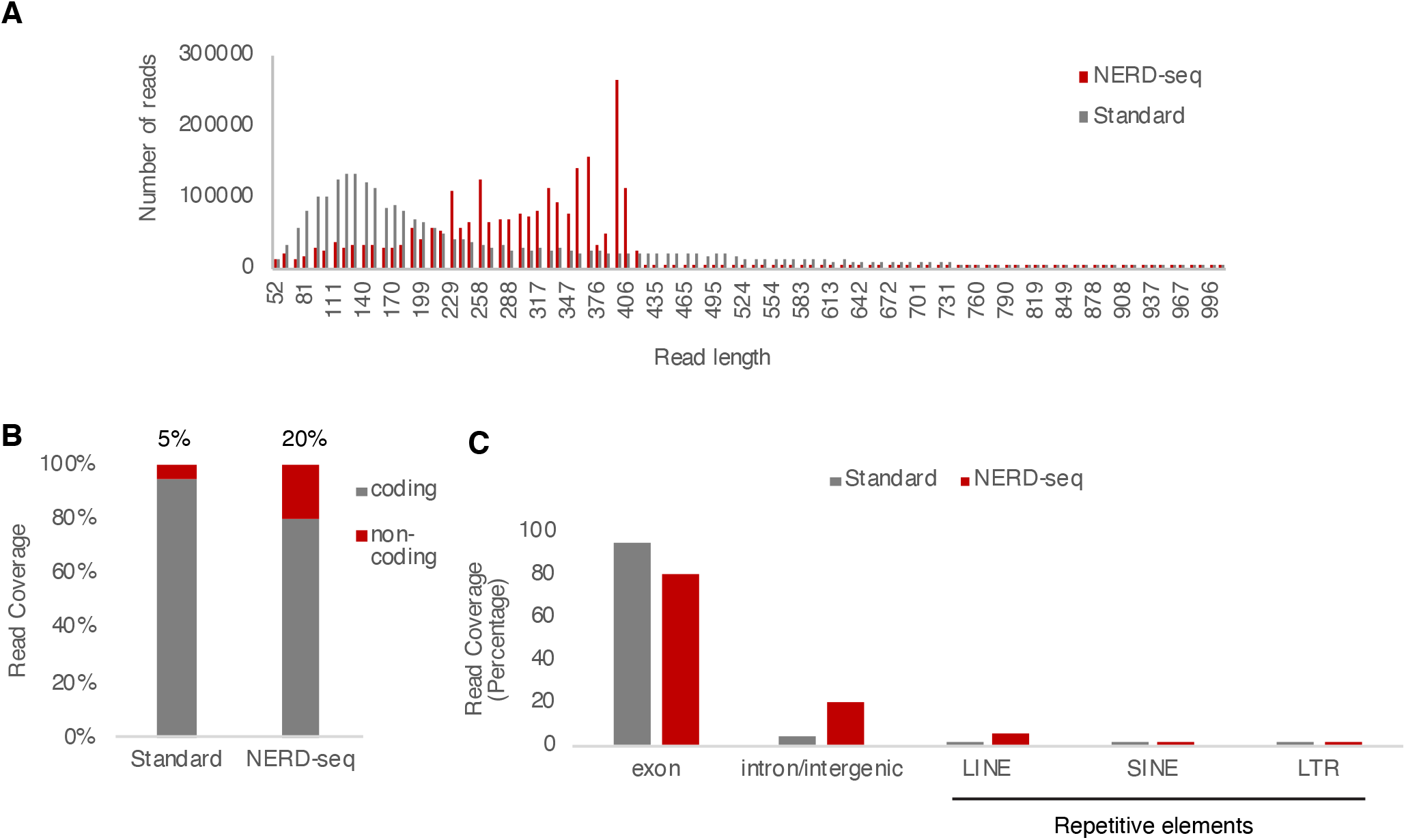
NERD-seq enables the generation of reads with higher coverage for the non-coding genome. **(A)** Comparison of read length distribution between the standard direct RNA-seq and the NERD-seq approach. **(B)** Percentage of reads aligning to the coding (exons) and non-coding portion of the genome in standard and NERD-seq approaches. **(C)** Coverage (percentage) across various coding and non-coding genomic elements for standard and NERD-seq reads.

### NERD-seq enables the generation of reads with higher coverage for the non-coding genome, while still detecting mRNAs and poly(A) ncRNAs

To assess the potential of NERD-seq we selected total RNA from mouse hippocampus, a tissue that is very active at the transcriptome and epitranscriptome level(61–63), and performed both NERD-seq and standard direct RNA-seq on the same total RNA from this tissue (Figure 1C). To our knowledge, there is only one study that has performed direct RNA-seq in neural tissues (whole brain)(64), and data from this study have been used as an external dataset for the validation of our findings (see supplement). While in that study Sessegolo et al. used the first developed platform by ONT, called MinION, we used here our in-house ONT PromethION platform, that has approx. a 10-time more yield compared to MinION. This enabled for a first time a more in-depth Nanopore sequencing in a neural mouse tissue, and the first direct RNA-seq in hippocampus. Both the standard approach and NERD-seq produced very high and comparable yields at 3.42 GigaBases (GB) and 3.48 GB, respectively. Consistent with the protocol for NERD-seq that ensures a higher representation of short RNAs, this similar yield in the overall number of sequenced bases translates to different read numbers, (3.6M for standard compared vs. 9.2M for NERD-seq). In other words, NERD-seq is able to produce more reads without this affecting its overall base yield. Moreover, NERD-seq achieved an even larger N50 number than the standard protocol. Read N50 refers to a value where half of the data is contained within reads with alignable lengths greater than this. N50 for NERD-seq was 1.84 KB compared to 1.16KB for the standard approach. Consistent with the high-temperature first-strand synthesis approach, these results confirmed the potential of NERD-seq to increase the average length of produced reads at the range around 400nt, to which many ncRNAs belong. Indeed, as shown in Figure 2A, the center of the NERD-seq read length distribution (red distribution) is skewed towards reads of higher average length (around 400nt). These results show that the inclusion of short and long non-poly(A) RNAs does not affect the overall yield metrics but does enrich for reads with length towards 400nt.

Then, we questioned how NERD-seq reads compare to those produced by the standard approach with regards to their alignment to some of the most prevalent coding and non-coding genome elements. To this end, we calculated the percentage of the sequenced reads overlapping (>80% of their length) to coding genes (known exons in mm10) or to regions outside of them (introns, intergenic regions). As shown in Figure 2B, standard direct-RNA-seq reads predominantly (95% of the total reads) represent coding regions and particularly known exons, confirming that the standard approach is primarily tailored towards detecting and studying mRNAs and protein-coding genes. In contrast, this percentage falls to 80% in the case of NERD-seq, with the portion of reads coming from non-coding regions climbing from 5% to 20%. Finally, LINE elements are among the major non-coding elements that are overrepresented in NERD-seq compared to the standard approach (Figure 2C).

Despite the increase of the percentage of reads representing introns and intergenic regions in NERD-seq, reads originating from exons still constitute 80% of overall reads, suggesting that NERD-seq remains efficient in detecting mRNAs. To test this, we constructed metagene models of all known genes and compared the relative read density around their transcription site (i.e. read numbers normalized to the total number of sample reads and elements that constitute the metagene). The generation of distribution plots of relative read densities allows comparisons of read coverage among different samples for the same set of genomic elements that construct the metagene model. As shown in Figure 3A, NERD-seq has been able to detect mRNAs as documented by the peak in the read distribution directly downstream of Transcription Start Site (TSS). Consistent with the higher enrichment in mRNAs in standard direct RNA-seq vs. NERD-seq mentioned above, the relative height of the peak of the distribution at TSS is higher in the standard than in NERD-seq (Kolmogorov–Smirnov test (KS test) < 0.05). However, a closer look to the characteristics of these distributions reveals that, despite being lower at the TSS, the NERD-seq distribution consists of proportionally longer transcripts that extend further downstream of TSS. This is more obvious when we scale both distributions (Supplementary Figure S1), revealing that reads from mRNAs detected by NERD-seq around TSS indeed tend in average to be longer than in the standard approach.

**Figure 3.**
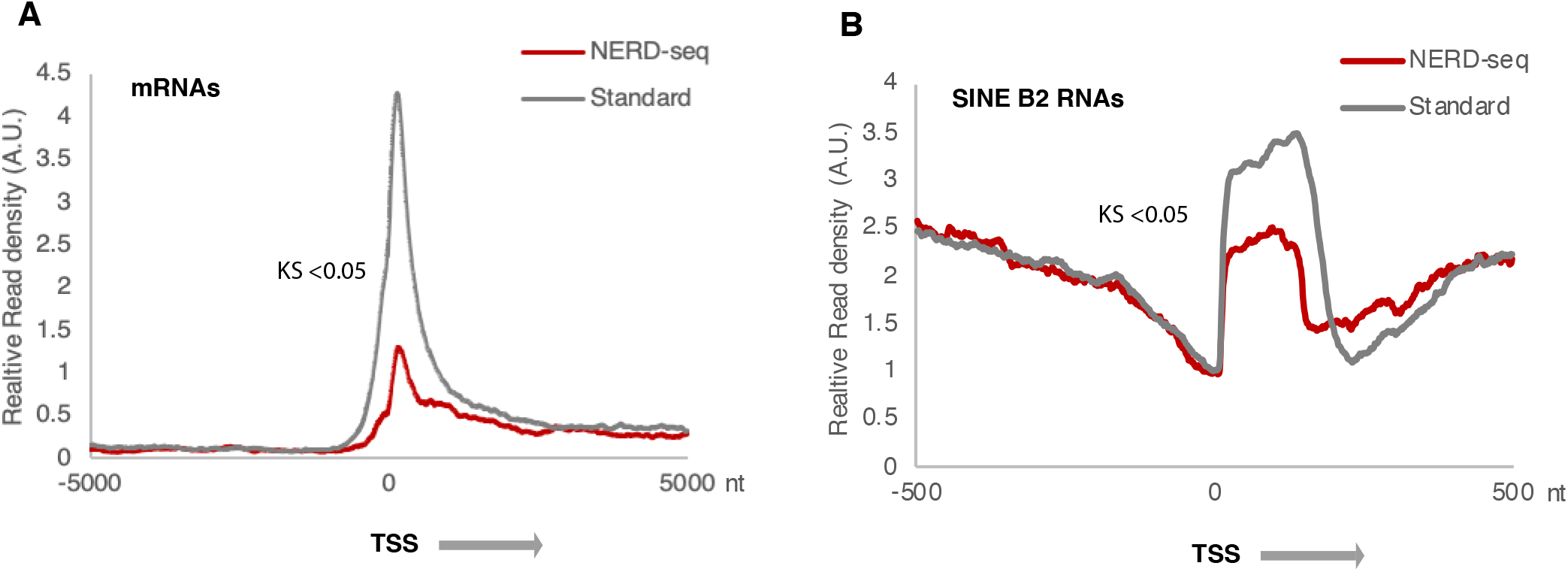
NERD-seq can detect mRNAs and poly(A) ncRNAs. **(A)** Relative read density distribution around the Transcription Start Site (TSS) of known genes for standard and NERD-seq. A metagene model has been constructed by aligning all known genes at their Transcription Start Site (as estimated by the use of Eponine at mm10(51)). Distances at X axis correspond to absolute distance (in nucleotides; nt) 500nt upstream (left) and downstream (right) from TSS. The arrow next to TSS depicts the direction of transcription. Read density is calculated by dividing the number of reads aligning to each position, divided to the total number of reads and elements (genes) that construct the metagene. KS: Kolmogorov–Smirnov test (KS) < 0.05 for the comparison between the two distributions. **(B)** Relative read density distribution around the Transcription Start Site (TSS) of SINE B2 RNAs (repeat masker mm10) for standard and NERD-seq.

Next, we asked whether NERD-seq can detect also ncRNAs that are known to be polyadenylated. To test this, we checked for the expression of ncRNAs generated by one of the most frequent SINE elements, the B2 element, which has been described to be polyadenylated(65,66). As shown in Figure 3B, as in the case of mRNAs, standard direct RNA-seq reads are more enriched in B2 RNAs, however, NERD-seq can still detect full-length B2 RNAs at sufficient levels.

Overall, our findings show that NERD-seq can expand coverage of non-coding genome while still being able to detect poly(A) transcripts detected by the standard protocol.

### NERD-seq can efficiently detect various classes of short non-poly(A) ncRNAs in contrast to the standard approach

We then questioned whether NERD-seq is able to detect known classes of non-poly(A) ncRNAs, which has been also one of our major motives for developing this method. To this end, we have first generated the relative read density distribution plots around the TSS of four classes of ncRNAs: snoRNAs, snRNAs, scRNAs and srpRNAs. As mentioned above, for validating our results, in order to exclude any lab-specific technical systematic errors, in addition to the data generated by us through the standard approach for the same RNA pool, we have also employed external direct RNA-seq data generated from the same organ (brain) and the same standard approach(64) for the comparison with our NERD-seq data (presented in Supplementary Figures S2 and S3).

As shown in Figure 4, for three out of four of these classes, snoRNAs, snRNAs and srpRNAs, the standard approach can hardly detect any of them, while it also significantly underperforms in the case of scRNAs (KS < 0.05) compared to NERD-seq. In contrast, NERD-seq produces robust distributions for all four classes (see also Supplementary Figure S2 for a comparison with external data). Then, we asked whether NERD-seq can detect another important class of non-poly(A) ncRNAs, tRNAs, that have been shown to be very frequent target of RNA modifications. As shown in Figure 5A, NERD-seq produces a robust distribution around the TSS of these RNAs (see also Supplementary Figure S3 for a comparison with external data). Interestingly though, as in the case of scRNAs, the standard approach can still detect some of them, though almost three-fold less than NERD-seq. Although tRNAs are known to be non-polyadenylated, it appears that they can still be detected by a poly(A)-selecting approach, such as the standard direct RNA-seq, presumably when they are marked with poly(A)s during degradation(67). As discussed below, this finding denotes the importance of an RNA-seq approach that can detect the non-poly(A) RNAs, as those may represent an entirely different biological context compared to the ones marked for degradation.

**Figure 4.**
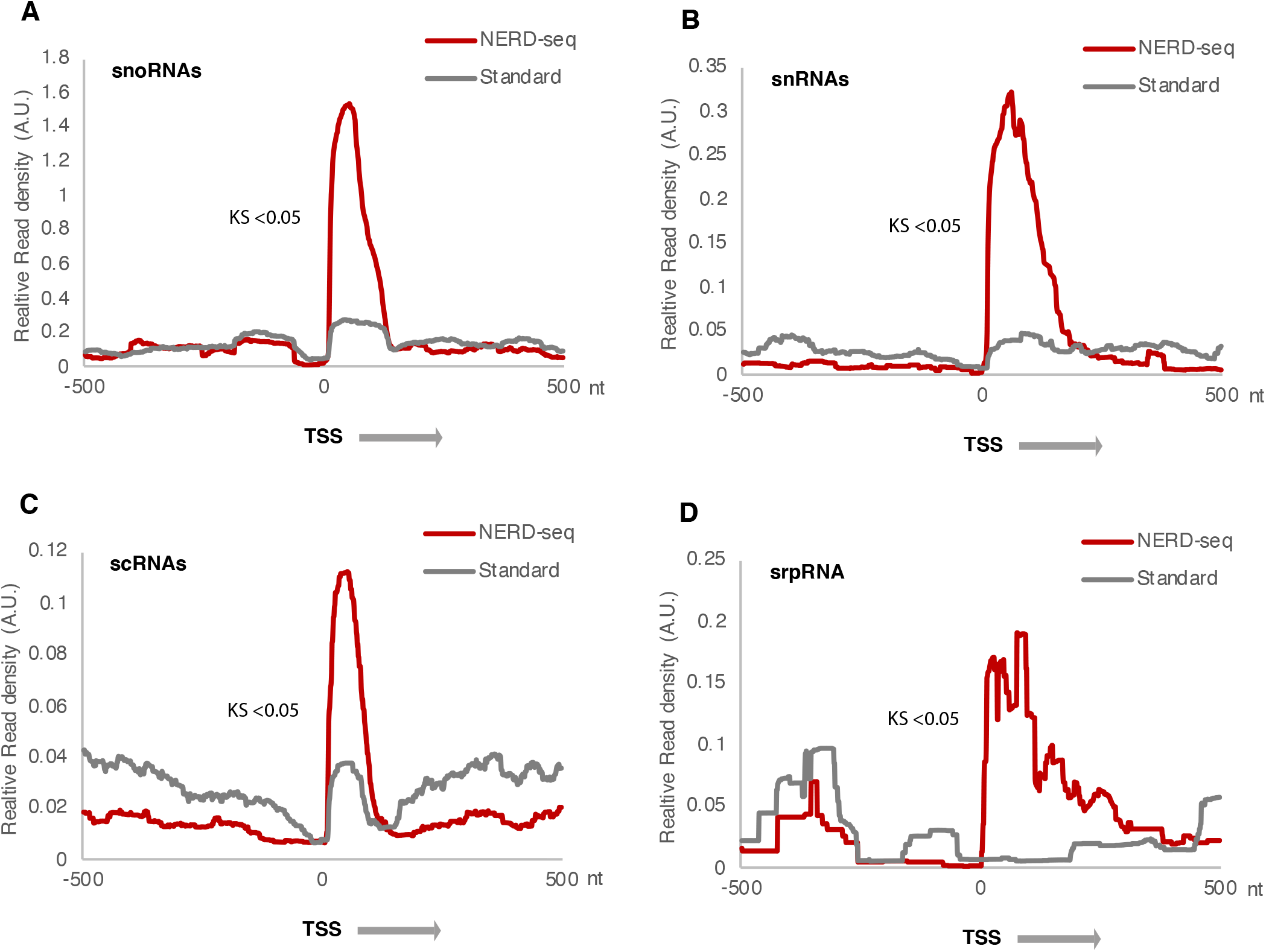
NERD-seq can detect snoRNAs, snRNAs, scRNAs and srpRNAs. **(A)** Relative read density distribution around the Transcription Start Site (TSS) of snoRNAs for standard and NERD-seq. X, Y axis and KS-test as in Figure 3. **(B)** Relative read density distribution around the Transcription Start Site (TSS) of snRNAs. **(C)** Relative read density distribution around the Transcription Start Site (TSS) of scRNAs. **(D)** Relative read density distribution around the Transcription Start Site (TSS) of srpRNA.

**Figure 5.**
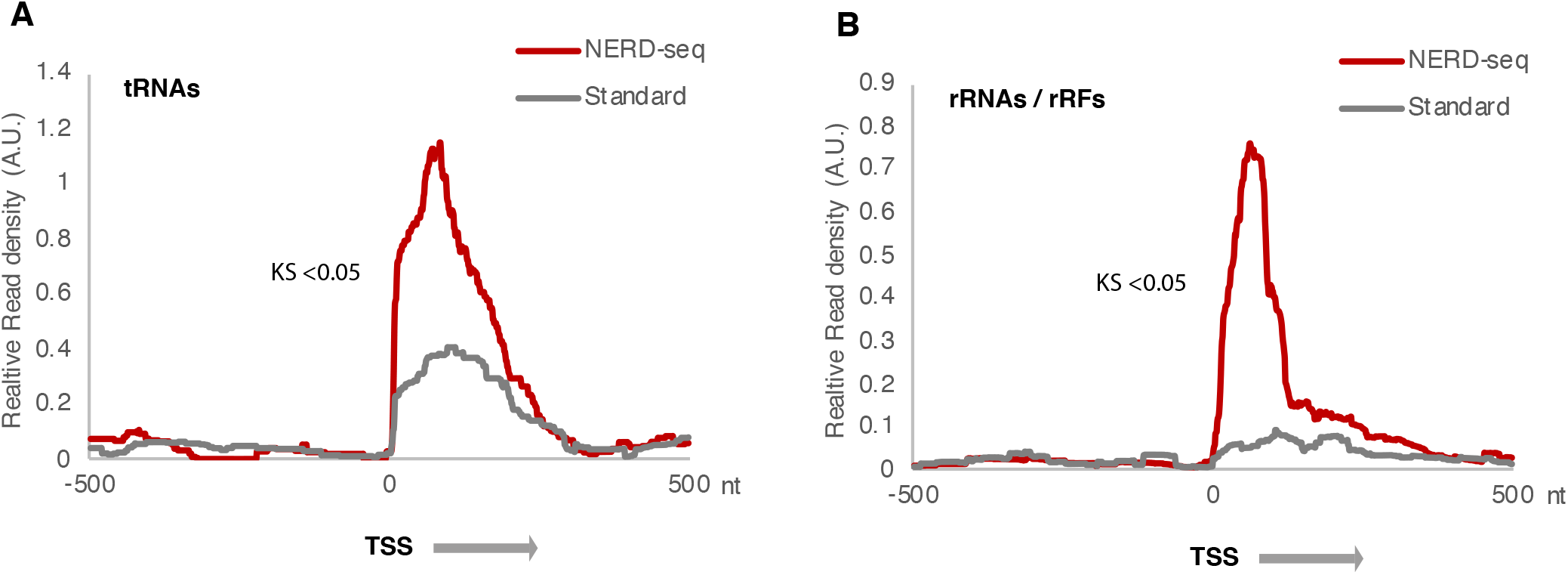
NERD-seq can detect tRNAs and ribosomal RNA fragments (rRFs). **(A)** Relative read density distribution around the Transcription Start Site (TSS) of known tRNAs (mm10) for standard and NERD-seq. X, Y axis and KS-test as in Figure 3. **(B)** Relative read density distribution around the Transcription Start Site (TSS) of rRNAs (mm10).

Finally, we examined the level of rRNA contamination in our data. As in the case of tRNAs, rRNAs are also marked with poly(A) tails for degradation, so even the standard approach is expected to detect some long rRNAs as shown in Fig.5B, which also applies to NERD-seq. Interestingly, in addition to the background rRNA detected in the standard approach, NERD-seq distribution around the rRNA TSS depicts a strong peak approx. 120-140nt wide, suggesting the existence of short rRNA fragments generated from this position that are missed by the standard approach (see Supplementary Figure S3 for a comparison with external data). Such rRNA fragments called rRFs, have been described as being capable of modulating rRNA transcription and function(68). As well, these fragments are important to cell survival and proliferation(69) in a sex, tissue, and population specific manner(70).

These results show that NERD-seq can efficiently expand direct RNA-seq capabilities to detect multiple classes of short non-poly(A) ncRNAs that may be missed by standard RNA-seq.

### NERD-seq allows for the detection of LINE 1 produced ncRNAs

As shown in Figure 2C, reads mapping to LINE elements are overrepresented in NERD-seq. ncRNAs from LINEs have been described in the past to be important for preventing neurodegeneration with its interaction with homeoprotein b in dopaminergic neurons(71). Also, they function to aid SINE RNAs in retrotransposition(72,73). LINEs span genomic regions around 5KB or more. We wanted to identify hotspots across the LINE elements producing the RNAs detected by NERD-seq. To this end, we mapped reads generated by both the NERD-seq and standard approach across the LINE metagene, which was constructed by aligning all known LINE elements in mm10 at their start site (Figures 6A and B, respectively). As shown in these figures, consistent with our findings in Figure 2C, LINE elements in NERD-seq data present multiple locations mapping ncRNAs compared to the standard approach. We then attempted to pinpoint the exact LINE elements producing these ncRNAs. To this end, we compared the read coverage of NERD-seq reads across all the major LINE elements listed in the UCSC mm10 repeat masker annotation track and presented at Fig.6C the three top-ranking ones based on read coverage numbers. As shown in this figure, NERD-seq read coverage of LINE elements comes predominantly from the L1 family, which far outweighs the following two families. This is confirmed by plotting NERD-seq reads at the respective L1 metagene (Figure 6D) and comparing this plot with that of other LINE families, such as L2 (Figure 6E). These results reveal the ability of NERD-seq to detect ncRNAs derived from LINE elements.

**Figure 6.**
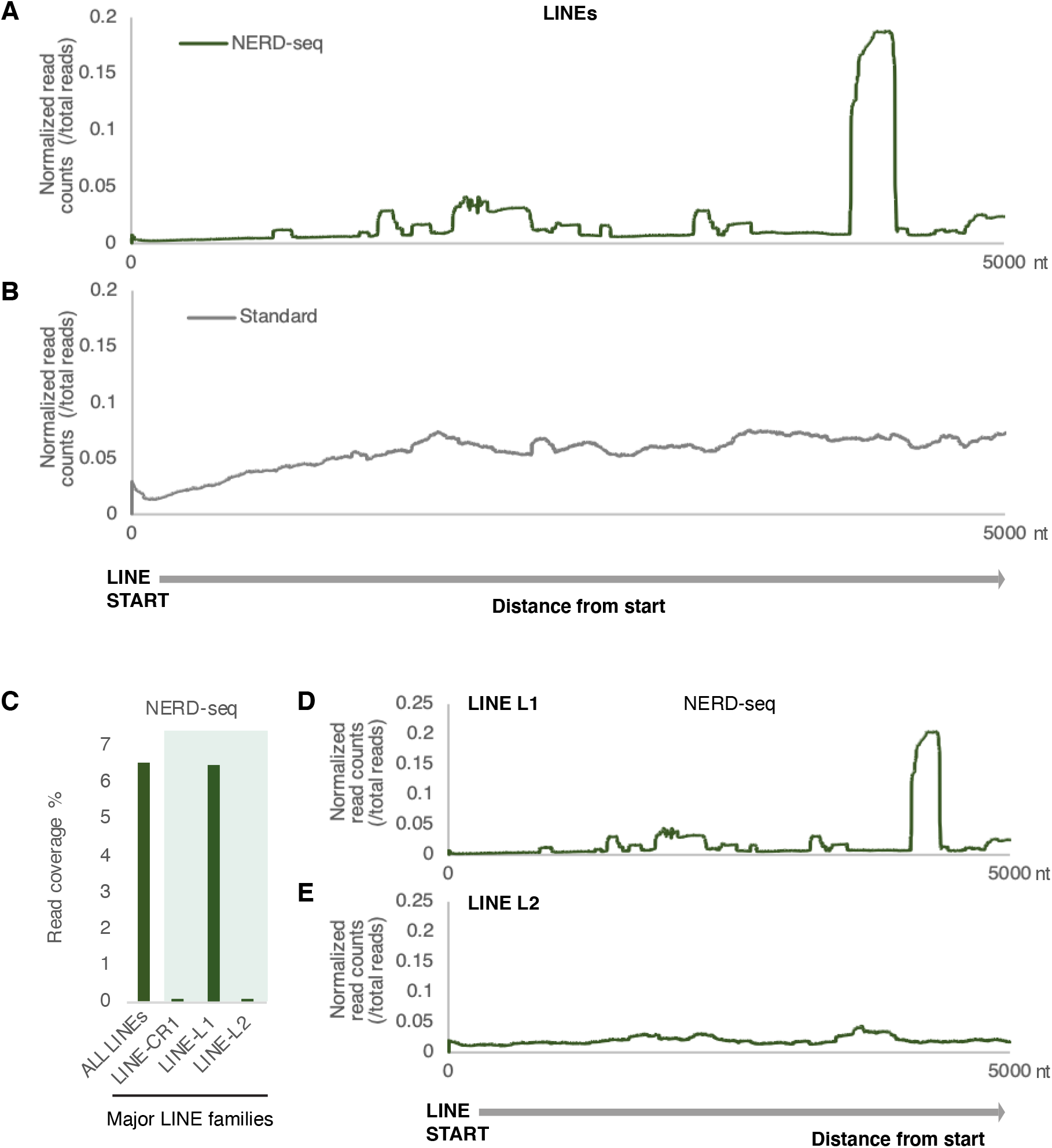
NERD-seq detects LINE L1 associated ncRNAs. **(A)** Normalized counts of reads from NERD-seq mapped across the first 5000nt of a metagene constructed by all known LINE elements (repeat masker UCSC mm10). The metagene model has been constructed by aligning all known LINEs at their start site (base #1 in each element’s DNA sequence). Distances at X axis correspond to absolute distance (in nucleotides; nt) 5000nt downstream from the start. The arrow below the graph corresponds to the sense direction of the elements. Normalized read counts per position are calculated by dividing the number of reads aligning to each position to the total number of reads. **(B)** Same as in (A) but for standard direct RNA-seq reads. **(C)** Coverage (percentage) across all LINEs and across the three LINE subfamilies for NERD-seq reads. The three LINE subfamilies with most aligned reads have been selected and are depicted here. **(D)** Normalized counts of reads from NERD-seq mapped across the first 5000nt of a metagene constructed as in (A) but by all known LINE L1 family elements. **(E)** Normalized counts of reads from NERD-seq mapped across the first 5000nt of a metagene constructed as in (A) but by all known LINE L2 family elements.

## DISCUSSION

Expanding our understanding of the epitranscriptome and its constituting RNA modifications, and why and where RNA edits occur, is expected to have profound influence on our understanding of how cells use RNA in different signalling and functional contexts(74,75), and how cells recognize internally transcribed RNAs compared to dysfunctional RNAs and RNA-related threats. The Nanopore platform and direct RNA-seq have opened new avenues for such studies. However, the standard direct RNA-seq approach prevents us from leveraging the full potential of this technology by excluding the detection of critical RNA classes that constitute some of the main targets of such modifications. Here, we present an approach that addresses these issues. NERD-seq can not only successfully detect mRNAs and poly(A) ncRNAs detected by the standard approach, but expands the detection of ncRNAs to all major classes of non-poly(A) ncRNAs. At the expense of reducing coverage of protein-coding RNAs for only 15% (decrease of 95% of overall reads to 80%), NERD-seq allows the successful detection of snoRNAs, snRNAs, scRNAs, srpRNAs and tRNAs. With the evolution of the Nanopore technology and the significant increase in the sequencing yield through ONT platforms such as PromethION, this small decrease in coverage of mRNAs can easily be tolerated. Moreover, while not amplifying full-length rRNAs, which is often undesirable, it can still detect rRFs that has recently proved to be critical in various biological contexts such as cell proliferation and survival(69) and rRNA transcription modulation(68).

For our comparison between NERD-seq and the standard protocol we chose hippocampal tissue due to its rich epitranscriptome variability throughout neurodevelopment and upon conditional treatment(61–63). Hippocampi exhibit high transcriptional activity and genome-environment interactions, while also being responsive to environmental and physiological changes(76–79). Additionally, it is a tissue that has received considerable attention for its role in memory formation and learning, and its early dysfunction in disorders such as Alzheimer’s and Parkinson’s disease and during cancer treatment(45,52,80–85).

Application of NERD-seq revealed similar metrics regarding base yield and a slightly improved N50, while generating almost three times the number of reads generated by the standard approach. This is a critical point, given that a significant increase in the number of RNA fragments to be sequenced could have reduced pore viability, motor protein fuel reserves and, thus, overall yield. An important observation that has arisen from our optimization experiments for this approach is the importance of adding random primers during the first-strand synthesis. Performing the first-strand synthesis at a higher temperature has undisputable advantages in resolving RNA structures that could impede the process. However, at this temperature RNA degradation is highly accelerated, which could result in a huge increase in the proportion of very short fragments (and overall of RNA fragments to be sequenced), which, as mentioned above, could be overwhelming for fuel reserves and detrimental for yield. Addition of random primers and a preliminary first-strand synthesis step at a lower temperature (50°C) before proceeding to higher temperature seems to prevent this and protect RNA from such degradation.

Our approach has enabled for the first time in-depth Nanopore PromethION sequencing in a neural mouse tissue, and the first direct RNA-seq in hippocampus ever with both the standard and our new NERD-seq approach. All classes that are detected by NERD-seq have been shown to bear critical roles in neural cells, and, thus, by expanding capabilities for studying their epitranscriptomes, NERD-seq will help further understand the mechanisms underlying this type of regulation. For example, snoRNAs have been shown to be key players in the RNA modification machinery but, until now, the standard approach does not allow to test for the impact of such changes on snoRNAs themselves in terms of potential self-regulatory loops. Moreover, marking of ncRNAs with poly(A) for degradation makes them the only ncRNAs in their class to be detected by the standard sequencing methods, adding a significant confounding factor, as any detected RNA modifications detected may be only connected with their degradation process and not with their other functions. NERD-seq can now identify those otherwise undetectable ncRNAs and increase the overall portfolio of RNAs

Nevertheless, this study leaves unanswered some important questions. For example, NERD-seq is able to detect more ncRNAs derived by LINEs compared to the standard approach, a finding that we have narrowed down to L1 elements, however, this finding is complicated by the large length of L1 elements. In particular, the identity of these L1-associated RNAs and whether they are related with previous ncRNAs described in LINEs, such as clusters of piRNAs, is unclear. Our attempt to associate the location of these ncRNAs with known locations of piRNAs was inconclusive, partially due to the large number of piRNA locations and their wide genomic distribution. Unfortunately, Nanopore sequencing does not allow for accurate basecalling of small RNAs such as piRNAs, which means that we cannot use the current datasets to search for a correlation between piRNA locations and that of our identified L1-associated ncRNAs. Application of Illumina sequencing has been neither conclusive due to the repetitive nature of these elements. Elucidating the identity of these ncRNAs lies beyond the scope of the current study, and future studies should elucidate the nature of these RNAs.

Overall, based on our findings, NERD-seq presents a simple but powerful approach for transcriptome and epitranscriptome analysis that expands our ability to exploit in full the potential of the Nanopore sequencing technology.

## ACKNOWLEDGEMENTS

This work has been supported by an Explorations Grant # 201700011 to AZ from Alberta Innovates and the Alberta Prion Research Institute, a Grant # 201900003 to AZ from the Alzheimer Society of Alberta and Northwest Territories and the Alberta Prion Research Institute, a Discovery Grant # RGPIN-2018-05955 to AZ from NSERC, the BioNet Alberta grant to AZ from Genome Canada, a AMR One Health grant by the Government of Alberta to AZ and a Compute Canada Resource Allocation Grant to AZ. AZ is supported by the Canada Research Chairs Program and the Canada Foundation for Innovation and is a former EMBO and DFG long-term fellow. YC, LS and LM are supported by an Alberta Innovates fellowship. LS and TH are supported by the AMR One Health grant by the Government of Alberta. We are grateful to Dr. Angeliki Pantazi for extensively reviewing, editing and commenting on the manuscript.

## AUTHOR CONTRIBUTIONS

LS: testing of the next-generation-sequencing, library construction, sequencing, writing of the manuscript; YC: Bioinformatics analysis, statistical analysis, establishment and testing of the analysis pipelines, data visualization; BG: testing of the next-generation-sequencing, library construction, sequencing; LM: Bioinformatics analysis, statistical analysis, establishment and testing of the analysis pipelines, data visualization; TH: testing of the next-generation-sequencing pipeline; MS: testing of the base calling pipeline; MM: data interpretation; AZ: conception and design, establishment and testing of data generation and analysis pipelines, bioinformatics analysis, data interpretation, data visualization and writing of the manuscript, overall supervision.

## CONFLICT OF INTEREST

The authors declare no conflict of interest.

## REFERENCES

1. Emrich, S.J., Barbazuk, W.B., Li, L. and Schnable, P.S. (2007) Gene discovery and annotation using LCM-454 transcriptome sequencing. Genome Res, 17, 69–73.

2. Lister, R., O’Malley, R.C., Tonti-Filippini, J., Gregory, B.D., Berry, C.C., Millar, A.H. and Ecker, J.R. (2008) Highly integrated single-base resolution maps of the epigenome in Arabidopsis. Cell, 133, 523–536.

3. Wang, Z., Gerstein, M. and Snyder, M. (2009) RNA-Seq: a revolutionary tool for transcriptomics. Nat Rev Genet, 10, 57–63.

4. Wagh, K., Ishikawa, M., Garcia, D.A., Stavreva, D.A., Upadhyaya, A. and Hager, G.L. (2021) Mechanical Regulation of Transcription: Recent Advances. Trends Cell Biol.

5. Nam, K.N., Mounier, A., Wolfe, C.M., Fitz, N.F., Carter, A.Y., Castranio, E.L., Kamboh, H.I., Reeves, V.L., Wang, J., Han, X. et al. (2017) Effect of high fat diet on phenotype, brain transcriptome and lipidome in Alzheimer’s model mice. Sci Rep, 7, 4307.

6. Dal Santo, S., Zenoni, S., Sandri, M., De Lorenzis, G., Magris, G., De Paoli, E., Di Gaspero, G., Del Fabbro, C., Morgante, M., Brancadoro, L. et al. (2018) Grapevine field experiments reveal the contribution of genotype, the influence of environment and the effect of their interaction (GxE) on the berry transcriptome. Plant J, 93, 1143–1159.

7. Wagner, P.J., Park, H.R., Wang, Z., Kirchner, R., Wei, Y., Su, L., Stanfield, K., Guilarte, T.R., Wright, R.O., Christiani, D.C. et al. (2017) In Vitro Effects of Lead on Gene Expression in Neural Stem Cells and Associations between Up-regulated Genes and Cognitive Scores in Children. Environ Health Perspect, 125, 721–729.

8. Cieslik, M. and Chinnaiyan, A.M. (2018) Cancer transcriptome profiling at the juncture of clinical translation. Nat Rev Genet, 19, 93–109.

9. Twine, N.A., Janitz, K., Wilkins, M.R. and Janitz, M. (2011) Whole transcriptome sequencing reveals gene expression and splicing differences in brain regions affected by Alzheimer’s disease. PLoS One, 6, e16266.

10. Vera, J.C., Wheat, C.W., Fescemyer, H.W., Frilander, M.J., Crawford, D.L., Hanski, I. and Marden, J.H. (2008) Rapid transcriptome characterization for a nonmodel organism using 454 pyrosequencing. Mol Ecol, 17, 1636–1647.

11. Kumar, R., Ichihashi, Y., Kimura, S., Chitwood, D.H., Headland, L.R., Peng, J., Maloof, J.N. and Sinha, N.R. (2012) A High-Throughput Method for Illumina RNA-Seq Library Preparation. Front Plant Sci, 3, 202.

12. Syddall, C.M., Reynard, L.N., Young, D.A. and Loughlin, J. (2013) The identification of trans-acting factors that regulate the expression of GDF5 via the osteoarthritis susceptibility SNP rs143383. PLoS Genet, 9, e1003557.

13. Tilgner, H., Jahanbani, F., Gupta, I., Collier, P., Wei, E., Rasmussen, M. and Snyder, M. (2018) Microfluidic isoform sequencing shows widespread splicing coordination in the human transcriptome. Genome Res, 28, 231–242.

14. Bolisetty, M.T., Rajadinakaran, G. and Graveley, B.R. (2015) Determining exon connectivity in complex mRNAs by nanopore sequencing. Genome Biol, 16, 204.

15. Kennedy, K., Hall, M.W., Lynch, M.D., Moreno-Hagelsieb, G. and Neufeld, J.D. (2014) Evaluating bias of illumina-based bacterial 16S rRNA gene profiles. Appl Environ Microbiol, 80, 5717–5722.

16. Lahens, N.F., Kavakli, I.H., Zhang, R., Hayer, K., Black, M.B., Dueck, H., Pizarro, A., Kim, J., Irizarry, R., Thomas, R.S. et al. (2014) IVT-seq reveals extreme bias in RNA sequencing. Genome Biol, 15, R86.

17. Ozsolak, F. and Milos, P.M. (2011) RNA sequencing: advances, challenges and opportunities. Nat Rev Genet, 12, 87–98.

18. Karst, S.M., Ziels, R.M., Kirkegaard, R.H., Sorensen, E.A., McDonald, D., Zhu, Q., Knight, R. and Albertsen, M. (2021) High-accuracy long-read amplicon sequences using unique molecular identifiers with Nanopore or PacBio sequencing. Nat Methods, 18, 165–169.

19. Zhao, L., Zhang, H., Kohnen, M.V., Prasad, K., Gu, L. and Reddy, A.S.N. (2019) Analysis of Transcriptome and Epitranscriptome in Plants Using PacBio Iso-Seq and Nanopore-Based Direct RNA Sequencing. Front Genet, 10, 253.

20. Chen, S.Y., Deng, F., Jia, X., Li, C. and Lai, S.J. (2017) A transcriptome atlas of rabbit revealed by PacBio single-molecule long-read sequencing. Sci Rep, 7, 7648.

21. Gradel, C., Terrazos Miani, M.A., Baumann, C., Barbani, M.T., Neuenschwander, S., Leib, S.L., Suter-Riniker, F. and Ramette, A. (2020) Whole-Genome Sequencing of Human Enteroviruses from Clinical Samples by Nanopore Direct RNA Sequencing. Viruses, 12.

22. Weirather, J.L., de Cesare, M., Wang, Y., Piazza, P., Sebastiano, V., Wang, X.J., Buck, D. and Au, K.F. (2017) Comprehensive comparison of Pacific Biosciences and Oxford Nanopore Technologies and their applications to transcriptome analysis. F1000Res, 6, 100.

23. Workman, R.E., Tang, A.D., Tang, P.S., Jain, M., Tyson, J.R., Razaghi, R., Zuzarte, P.C., Gilpatrick, T., Payne, A., Quick, J. et al. (2019) Nanopore native RNA sequencing of a human poly(A) transcriptome. Nat Methods, 16, 1297–1305.

24. Soneson, C., Yao, Y., Bratus-Neuenschwander, A., Patrignani, A., Robinson, M.D. and Hussain, S. (2019) A comprehensive examination of Nanopore native RNA sequencing for characterization of complex transcriptomes. Nat Commun, 10, 3359.

25. Garalde, D.R., Snell, E.A., Jachimowicz, D., Sipos, B., Lloyd, J.H., Bruce, M., Pantic, N., Admassu, T., James, P., Warland, A. et al. (2018) Highly parallel direct RNA sequencing on an array of nanopores. Nat Methods, 15, 201–206.

26. Price, A.M., Hayer, K.E., McIntyre, A.B.R., Gokhale, N.S., Abebe, J.S., Della Fera, A.N., Mason, C.E., Horner, S.M., Wilson, A.C., Depledge, D.P. et al. (2020) Direct RNA sequencing reveals m(6)A modifications on adenovirus RNA are necessary for efficient splicing. Nat Commun, 11, 6016.

27. Wang, L., Si, Y., Dedow, L.K., Shao, Y., Liu, P. and Brutnell, T.P. (2011) A low-cost library construction protocol and data analysis pipeline for Illumina-based strand-specific multiplex RNA-seq. PLoS One, 6, e26426.

28. Smith, A.M., Jain, M., Mulroney, L., Garalde, D.R. and Akeson, M. (2019) Reading canonical and modified nucleobases in 16S ribosomal RNA using nanopore native RNA sequencing. PLoS One, 14, e0216709.

29. Ontiveros, R.J., Stoute, J. and Liu, K.F. (2019) The chemical diversity of RNA modifications. Biochem J, 476, 1227–1245.

30. Nishikura, K. (2010) Functions and regulation of RNA editing by ADAR deaminases. Annu Rev Biochem, 79, 321–349.

31. Bass, B.L. (2002) RNA editing by adenosine deaminases that act on RNA. Annu Rev Biochem, 71, 817–846.

32. Roth, S.H., Levanon, E.Y. and Eisenberg, E. (2019) Genome-wide quantification of ADAR adenosine-to-inosine RNA editing activity. Nat Methods, 16, 1131–1138.

33. Dunin-Horkawicz, S., Czerwoniec, A., Gajda, M.J., Feder, M., Grosjean, H. and Bujnicki, J.M. (2006) MODOMICS: a database of RNA modification pathways. Nucleic Acids Res, 34, D145–149.

34. Cantara, W.A., Crain, P.F., Rozenski, J., McCloskey, J.A., Harris, K.A., Zhang, X., Vendeix, F.A., Fabris, D. and Agris, P.F. (2011) The RNA Modification Database, RNAMDB: 2011 update. Nucleic Acids Res, 39, D195–201.

35. Schwartz, S., Bernstein, D.A., Mumbach, M.R., Jovanovic, M., Herbst, R.H., Leon-Ricardo, B.X., Engreitz, J.M., Guttman, M., Satija, R., Lander, E.S. et al. (2014) Transcriptome-wide mapping reveals widespread dynamic-regulated pseudouridylation of ncRNA and mRNA. Cell, 159, 148–162.

36. Crooke, P.S., 3rd, Tossberg, J.T., Porter, K.P. and Aune, T.M. (2021) Cutting Edge: Reduced Adenosine-to-Inosine Editing of Endogenous Alu RNAs in Severe COVID-19 Disease. J Immunol, 206, 1691–1696.

37. Tossberg, J.T., Heinrich, R.M., Farley, V.M., Crooke, P.S., 3rd and Aune, T.M. (2020) Adenosine-to-Inosine RNA Editing of Alu Double-Stranded (ds)RNAs Is Markedly Decreased in Multiple Sclerosis and Unedited Alu dsRNAs Are Potent Activators of Proinflammatory Transcriptional Responses. J Immunol, 205, 2606–2617.

38. Di Timoteo, G., Dattilo, D., Centron-Broco, A., Colantoni, A., Guarnacci, M., Rossi, F., Incarnato, D., Oliviero, S., Fatica, A., Morlando, M. et al. (2020) Modulation of circRNA Metabolism by m(6)A Modification. Cell Rep, 31, 107641.

39. Ramos, J., Proven, M., Halvardson, J., Hagelskamp, F., Kuchinskaya, E., Phelan, B., Bell, R., Kellner, S.M., Feuk, L., Thuresson, A.C. et al. (2020) Identification and rescue of a tRNA wobble inosine deficiency causing intellectual disability disorder. RNA, 26, 1654–1666.

40. Lorenz, D.A., Sathe, S., Einstein, J.M. and Yeo, G.W. (2020) Direct RNA sequencing enables m(6)A detection in endogenous transcript isoforms at base-specific resolution. RNA, 26, 19–28.

41. Liu, H., Begik, O., Lucas, M.C., Ramirez, J.M., Mason, C.E., Wiener, D., Schwartz, S., Mattick, J.S., Smith, M.A. and Novoa, E.M. (2019) Accurate detection of m(6)A RNA modifications in native RNA sequences. Nat Commun, 10, 4079.

42. Gong, J., Liu, C., Liu, W., Xiang, Y., Diao, L., Guo, A.Y. and Han, L. (2017) LNCediting: a database for functional effects of RNA editing in lncRNAs. Nucleic Acids Res, 45, D79–D84.

43. Athanasiadis, A., Rich, A. and Maas, S. (2004) Widespread A-to-I RNA editing of Alu-containing mRNAs in the human transcriptome. PLoS Biol, 2, e391.

44. Kim, D.D., Kim, T.T., Walsh, T., Kobayashi, Y., Matise, T.C., Buyske, S. and Gabriel, A. (2004) Widespread RNA editing of embedded alu elements in the human transcriptome. Genome Res, 14, 1719–1725.

45. Cheng, Y., Saville, L., Gollen, B., Isaac, C., Belay, A., Mehla, J., Patel, K., Thakor, N., Mohajerani, M.H. and Zovoilis, A. (2020) Increased processing of SINE B2 ncRNAs unveils a novel type of transcriptome deregulation in amyloid beta neuropathology. Elife, 9.

46. Agis-Balboa, R.C., Arcos-Diaz, D., Wittnam, J., Govindarajan, N., Blom, K., Burkhardt, S., Haladyniak, U., Agbemenyah, H.Y., Zovoilis, A., Salinas-Riester, G. et al. (2011) A hippocampal insulin-growth factor 2 pathway regulates the extinction of fear memories. EMBO J, 30, 4071–4083.

47. Hernandez, A.J., Zovoilis, A., Cifuentes-Rojas, C., Han, L., Bujisic, B. and Lee, J.T. (2020) B2 and ALU retrotransposons are self-cleaving ribozymes whose activity is enhanced by EZH2. Proc Natl Acad Sci U S A, 117, 415–425.

48. Zovoilis, A., Cifuentes-Rojas, C., Chu, H.P., Hernandez, A.J. and Lee, J.T. (2016) Destabilization of B2 RNA by EZH2 Activates the Stress Response. Cell, 167, 1788–1802 e1713.

49. Li, H., Handsaker, B., Wysoker, A., Fennell, T., Ruan, J., Homer, N., Marth, G., Abecasis, G., Durbin, R. and Genome Project Data Processing, S. (2009) The Sequence Alignment/Map format and SAMtools. Bioinformatics, 25, 2078–2079.

50. Quinlan, A.R. and Hall, I.M. (2010) BEDTools: a flexible suite of utilities for comparing genomic features. Bioinformatics, 26, 841–842.

51. Down, T.A. and Hubbard, T.J. (2002) Computational detection and location of transcription start sites in mammalian genomic DNA. Genome Res, 12, 458–461.

52. Cheng, Y., Saville, L., Gollen, B., Veronesi, A.A., Mohajerani, M., Joseph, J.T. and Zovoilis, A. (2021) Increased Alu RNA processing in Alzheimer brains is linked to gene expression changes. EMBO Rep, e52255.

53. Mraz, M., Malinova, K., Mayer, J. and Pospisilova, S. (2009) MicroRNA isolation and stability in stored RNA samples. Biochem Biophys Res Commun, 390, 1–4.

54. Wilson, B.D., Eisenstein, M. and Soh, H.T. (2019) High-Fidelity Nanopore Sequencing of Ultra-Short DNA Targets. Anal Chem, 91, 6783–6789.

55. Kono, N. and Arakawa, K. (2019) Nanopore sequencing: Review of potential applications in functional genomics. Dev Growth Differ, 61, 316–326.

56. Boivin, V., Deschamps-Francoeur, G., Couture, S., Nottingham, R.M., Bouchard-Bourelle, P., Lambowitz, A.M., Scott, M.S. and Abou-Elela, S. (2018) Simultaneous sequencing of coding and noncoding RNA reveals a human transcriptome dominated by a small number of highly expressed noncoding genes. RNA, 24, 950–965.

57. Chander, Y., Koelbl, J., Puckett, J., Moser, M.J., Klingele, A.J., Liles, M.R., Carrias, A., Mead, D.A. and Schoenfeld, T.W. (2014) A novel thermostable polymerase for RNA and DNA loop-mediated isothermal amplification (LAMP). Front Microbiol, 5, 395.

58. Aonuma, H., Iizuka-Shiota, I., Hoshina, T., Tajima, S., Kato, F., Hori, S., Saijo, M. and Kanuka, H. (2020) Detection and discrimination of multiple strains of Zika virus by reverse transcription-loop-mediated isothermal amplification. Trop Med Health, 48, 87.

59. Schoenfeld, T., Patterson, M., Richardson, P.M., Wommack, K.E., Young, M. and Mead, D. (2008) Assembly of viral metagenomes from yellowstone hot springs. Appl Environ Microbiol, 74, 4164–4174.

60. Moser, M.J., DiFrancesco, R.A., Gowda, K., Klingele, A.J., Sugar, D.R., Stocki, S., Mead, D.A. and Schoenfeld, T.W. (2012) Thermostable DNA polymerase from a viral metagenome is a potent RT-PCR enzyme. PLoS One, 7, e38371.

61. Li, L., Zang, L., Zhang, F., Chen, J., Shen, H., Shu, L., Liang, F., Feng, C., Chen, D., Tao, H. et al. (2017) Fat mass and obesity-associated (FTO) protein regulates adult neurogenesis. Hum Mol Genet, 26, 2398–2411.

62. Merkurjev, D., Hong, W.T., Iida, K., Oomoto, I., Goldie, B.J., Yamaguti, H., Ohara, T., Kawaguchi, S.Y., Hirano, T., Martin, K.C. et al. (2018) Synaptic N(6)-methyladenosine (m(6)A) epitranscriptome reveals functional partitioning of localized transcripts. Nat Neurosci, 21, 1004–1014.

63. Zhang, Z., Wang, M., Xie, D., Huang, Z., Zhang, L., Yang, Y., Ma, D., Li, W., Zhou, Q., Yang, Y.G. et al. (2018) METTL3-mediated N(6)-methyladenosine mRNA modification enhances long-term memory consolidation. Cell Res, 28, 1050–1061.

64. Sessegolo, C., Cruaud, C., Da Silva, C., Cologne, A., Dubarry, M., Derrien, T., Lacroix, V. and Aury, J.M. (2019) Transcriptome profiling of mouse samples using nanopore sequencing of cDNA and RNA molecules. Sci Rep, 9, 14908.

65. Borodulina, O.R. and Kramerov, D.A. (2008) Transcripts synthesized by RNA polymerase III can be polyadenylated in an AAUAAA-dependent manner. RNA, 14, 1865–1873.

66. Ustyantsev, I.G., Borodulina, O.R. and Kramerov, D.A. (2020) Identification of nucleotide sequences and some proteins involved in polyadenylation of RNA transcribed by Pol III from SINEs. RNA Biol, 1–14.

67. Toompuu, M., Tuomela, T., Laine, P., Paulin, L., Dufour, E. and Jacobs, H.T. (2018) Polyadenylation and degradation of structurally abnormal mitochondrial tRNAs in human cells. Nucleic Acids Res, 46, 5209–5226.

68. Zhu, C., Yan, Q., Weng, C., Hou, X., Mao, H., Liu, D., Feng, X. and Guang, S. (2018) Erroneous ribosomal RNAs promote the generation of antisense ribosomal siRNA. Proc Natl Acad Sci U S A, 115, 10082–10087.

69. Chen, Z., Sun, Y., Yang, X., Wu, Z., Guo, K., Niu, X., Wang, Q., Ruan, J., Bu, W. and Gao, S. (2017) Two featured series of rRNA-derived RNA fragments (rRFs) constitute a novel class of small RNAs. PLoS One, 12, e0176458.

70. Cherlin, T., Magee, R., Jing, Y., Pliatsika, V., Loher, P. and Rigoutsos, I. (2020) Ribosomal RNA fragmentation into short RNAs (rRFs) is modulated in a sex- and population of origin-specific manner. BMC Biol, 18, 38.

71. Blaudin de The, F.X., Rekaik, H., Peze-Heidsieck, E., Massiani-Beaudoin, O., Joshi, R.L., Fuchs, J. and Prochiantz, A. (2018) Engrailed homeoprotein blocks degeneration in adult dopaminergic neurons through LINE-1 repression. EMBO J, 37.

72. Elbarbary, R.A., Lucas, B.A. and Maquat, L.E. (2016) Retrotransposons as regulators of gene expression. Science, 351, aac7247.

73. Richardson, S.R., Doucet, A.J., Kopera, H.C., Moldovan, J.B., Garcia-Perez, J.L. and Moran, J.V. (2015) The Influence of LINE-1 and SINE Retrotransposons on Mammalian Genomes. Microbiol Spectr, 3, MDNA3-0061-2014.

74. Galeano, F., Tomaselli, S., Locatelli, F. and Gallo, A. (2012) A-to-I RNA editing: the “ADAR” side of human cancer. Semin Cell Dev Biol, 23, 244–250.

75. Kishi, H. (1986) [Recent trend in pathogenic bacteria of urinary tract infections--simple and complicated urinary tract infections]. Nihon Rinsho, 44, 2552–2557.

76. Fernandez-Albert, J., Lipinski, M., Lopez-Cascales, M.T., Rowley, M.J., Martin-Gonzalez, A.M., Del Blanco, B., Corces, V.G. and Barco, A. (2019) Immediate and deferred epigenomic signatures of in vivo neuronal activation in mouse hippocampus. Nat Neurosci, 22, 1718–1730.

77. Schulz, H., Ruppert, A.K., Herms, S., Wolf, C., Mirza-Schreiber, N., Stegle, O., Czamara, D., Forstner, A.J., Sivalingam, S., Schoch, S. et al. (2017) Genome-wide mapping of genetic determinants influencing DNA methylation and gene expression in human hippocampus. Nat Commun, 8, 1511.

78. Solvsten, C.A.E., de Paoli, F., Christensen, J.H. and Nielsen, A.L. (2018) Voluntary Physical Exercise Induces Expression and Epigenetic Remodeling of VegfA in the Rat Hippocampus. Mol Neurobiol, 55, 567–582.

79. Peleg, S., Sananbenesi, F., Zovoilis, A., Burkhardt, S., Bahari-Javan, S., Agis-Balboa, R.C., Cota, P., Wittnam, J.L., Gogol-Doering, A., Opitz, L. et al. (2010) Altered histone acetylation is associated with age-dependent memory impairment in mice. Science, 328, 753–756.

80. Lombardi, G., Crescioli, G., Cavedo, E., Lucenteforte, E., Casazza, G., Bellatorre, A.G., Lista, C., Costantino, G., Frisoni, G., Virgili, G. et al. (2020) Structural magnetic resonance imaging for the early diagnosis of dementia due to Alzheimer’s disease in people with mild cognitive impairment. Cochrane Database Syst Rev, 3, CD009628.

81. Hoozemans, J.J., van Haastert, E.S., Nijholt, D.A., Rozemuller, A.J., Eikelenboom, P. and Scheper, W. (2009) The unfolded protein response is activated in pretangle neurons in Alzheimer’s disease hippocampus. Am J Pathol, 174, 1241–1251.

82. Foo, H., Mak, E., Chander, R.J., Ng, A., Au, W.L., Sitoh, Y.Y., Tan, L.C. and Kandiah, N. (2017) Associations of hippocampal subfields in the progression of cognitive decline related to Parkinson’s disease. Neuroimage Clin, 14, 37–42.

83. Yang, M. and Moon, C. (2015) Effects of cancer therapy on hippocampus-related function. Neural Regen Res, 10, 1572–1573.

84. Pantazi, A. and Zovoilis, A. (2013) Vector-free methods for manipulating miRNA activity in vitro and in vivo. Methods Mol Biol, 936, 231–245.

85. Zovoilis, A., Agbemenyah, H.Y., Agis-Balboa, R.C., Stilling, R.M., Edbauer, D., Rao, P., Farinelli, L., Delalle, I., Schmitt, A., Falkai, P. et al. (2011) microRNA-34c is a novel target to treat dementias. EMBO J, 30, 4299–4308.

